# Spatial summation of individual cones in human color vision

**DOI:** 10.1101/521492

**Authors:** Brian P. Schmidt, Alexandra E. Boehm, William S. Tuten, Austin Roorda

## Abstract

The human retina contains three classes of cone photoreceptors each sensitive to different portions of the visual spectrum: long (L), medium (M) and short (S) wavelengths. Color information is computed by downstream neurons that compare relative activity across the three cone types. How cone signals are combined at a cellular scale has been more difficult to resolve. This is especially true near the fovea, where spectrally-opponent neurons in the parvocellular pathway draw excitatory input from a single cone and thus even the smallest stimulus will engage multiple color-signaling neurons. We used an adaptive optics microstimulator to target individual and pairs of cones with light. Consistent with prior work, we found that color percepts elicited from individual cones were predicted by their spectral sensitivity, although there was considerable variability even between cones within the same spectral class. The appearance of spots targeted at two cones were predicted by an average of their individual activations. However, two cones of the same subclass elicited percepts that were systematically more saturated than predicted by an average. Together, these observations suggest both spectral opponency and prior experience influence the appearance of small spots.

## Introduction

A central goal of vision science is to understand how signals from photoreceptors are transformed into sight and the limitations each stage of processing imposes on perception. Photoreceptors encode real-time information about the environment. However, the signals conveyed by individual neurons are noisy and ambiguous. One strategy for reducing uncertainty is to pool signals across multiple detectors. Under low light conditions, for example, the visual system combines signals from many hundreds of rod and cone photoreceptors in order to boost sensitivity [1]. One drawback of signal pooling is a loss in spatial resolution: acuity is reduced under low-light levels [2]. We studied the influence of spatial pooling on the color appearance of cone-targeted spots.

The role of spatial pooling on light detection has been well documented. In terms of energy (stimulus intensity multiplied by area), detection thresholds of uniform stimuli are independent of size below a certain critical value (Ricco’s area), known as the area of complete spatial summation. Within this area, summation is linear and threshold energy, i.e. the total number of quanta, remains constant. At larger stimulus sizes, which exceed the critical area, threshold energy increases. One mechanistic explanation of this phenomenon is that linearly summation occurs when a spot activates cones that feed into a common downstream neuron. When a spot exceeds the size of a single receptive field, the energy is split over multiple downstream neurons and each additional neuron introduces an independent noise source. Consequently, signal to noise ratio (SNR) is reduced and summation becomes sub-linear. The magnocellular pathway is often cited as a possible neural correlate of this phenomenon [3–5]. Recently, this idea was tested by Bruce [6], who measured summation on a cellular scale using an adaptive optics scanning laser ophthalmoscope (AOSLO). They stimulated individual and pairs of cones and measured detection thresholds. They found a fraction of stimulated cone pairs summed light sub-linearly, including some that were within Ricco’s area, which was consistent with activation of two channels carrying independent noise sources.

Cone signals are also integrated over space in color vision. Color-opponent neurons compare the relative activity between cones with different spectral sensitivities [7–9]. These spatial comparisons lead to a loss of resolution. Chromatic contrast sensitivity functions have a spatial frequency cutoff that is about a third of the achromatic subsystem [10, 11]. The location of a stimulus on the retina also influences color appearance. As a stimulus moves away from the center of gaze it tends to appear less saturated [12], but this effect can be offset by increasing the size of the stimulus [13, 14]. For instance, at five degrees eccentricity, Abramov et al. [13] found that a monochromatic 580 nm, 0.5 degrees stimulus appeared yellow and ~18% saturated. Saturation increased to ~48% when the diameter was doubled. Abramov et al. [13] argued that this interaction between size and eccentricity was a reflection of the underlying physiology. Both the diameter of receptive fields and the number of cones pooled increase as a function of eccentricity [15, 16] and this may explain why stimuli must be larger in the periphery to achieve equivalent saturation. As receptive fields grow larger, so too must chromatic stimuli in order to drive the same rate of neural activity. In the fovea, the same trend has been observed as stimulus size increases from 0.2 to 1 degree in diameter [17]. However, foveal color opponent neurons in the parvocellular pathway often draw excitatory input from single cones [15, 18], which are less than 0.01 degrees in diameter [19]. Thus, it is difficult to conclude from this work exactly how receptive field diameter factors into size dependent changes in saturation, since a 0.2 degree spot would excite many dozens of parvocellular neurons. Moreover, traditional psychophysical measurements introduce uncertainty about the exact position and distribution of light on the retina. The eye’s own natural movements coupled with optical blur make it difficult to directly relate perception to the activation of specific photoreceptors.

To overcome these challenges, we used an AOSLO to image, track and stimulate individual or pairs of cones [20]. A hue-scaling paradigm was used to quantify the color appearance of each stimulus. Previous work has recorded percepts elicited from individual cones and identified a few general trends. Firstly, the spectral sensitivity of a probed cone is an important factor governing the elicited percept [21–23]. Secondly, even cones with the same spectral sensitivity elicit different colors when probed with identical stimuli [21, 24]. These observations have been interpreted as evidence that, near the fovea, the visual system learns different information about each cone [21, 25]. How the visual system combines information from individual cones may depend on the qualities the brain has learned to associate with each signal.

Here we report that the color of small spots is influenced by the number and type of cones targeted. On average, when two L-cones were targeted they produced a more saturated red percept than predicted from their individual activations. In comparison, an L- and M-cone activated together tended to elicit desaturated percepts. Together, these observations implicate both spectral opponency and prior experience in the process of computing the color of very small spots.

## Methods

### Subjects

Three highly experienced subjects participated in the study. S10001 was a 34 year old male. S20075 was a 30 year old female. S20076 was a 31 year old male. All subjects had normal color vision (anomaloscope and Hardy-Rand-Rittler or Ishihara pseudoisochromatic plates) and were authors of the study. At the start of each session, cycloplegia and mydriasis were induced with drops of 1.0% tropicamide and 2.5% phenylephrine hydrochloride ophthalmic solution. All procedures were approved by the Institutional Review Board at the University of California Berkeley and adhered to the tenets of the Declaration of Helsinki. Informed consent was obtained from each subject before the experiments.

### AOSLO Microstimulator

A multi-wavelength adaptive optics scanning laser ophthalmoscope (AOSLO) was used to image and present stimuli to the retina. The AOSLO system [26–28] and the procedures for stimulating single cones have been described elsewhere in detail [20,21,23]. Briefly, light from a super-continuum laser (SuperK Extreme; NKT Photonics) was split into three channels with interference filters (Semrock): (1) A 940 nm channel was used to measure monochromatic aberrations. Light from this channel was collected into a wavefront sensor and that information was fed in real-time to a deformable mirror (DM97-08; ALPAO), which compensated for the measured aberrations. The resulting optical system was approximately diffraction limited [26]. (2) An 840 nm channel was used to image the retina. Light from this channel was collected into a photo-multiplier tube (H7422-50; Hamamatsu) via a confocal pinhole and rendered into a video stream. (3) A 543 nm channel was used for retinally-targeted stimulation. L- and M-cones are approximately equally sensitive to this wavelength [29].

Retinal tracking was performed following the procedures of Arathorn et al. [30]. Briefly, the 840 nm video stream was registered to a reference image with a strip based cross-correlation procedure, which output retinal coordinates. Those coordinates were used to drive an acousto-optic modulator (Brimrose Corp.), a high-speed optical switch, which modulated the 543 nm channel. When the raster scan passed over a cell of interest the switch opened and delivered a calibrated dose of light to the cell.

Chromatic aberration between the three channels was measured and corrected with established procedures [31]. The imaging and stimulation rasters subtended a 0.95 degree field at a sampling resolution of ~0.11 arcmin/pixel. The background in both experiments was white (CIE xy= 0.3, 0.32; 40 cd/m^2^). Subject’s heads were stabilized with a custom-fit bite bar. For additional details on single cone stimulation and the accuracy of this procedure, Meadway and Sincich [32] recently published a detailed model of light propagation and capture by cone photoreceptors in AOSLO systems.

### Cone classification

In two subjects (S10001 and S20076), cones were classified according to their spectral type (L, M, S) using densitomety. The details of that procedure have been described elsewhere [33, 34]. The accuracy of these measurements is approximately 95% [33]. In one subject, we were unable to collect images with sufficient SNR to reliably classify cones.

### Threshold measurements

Detection thresholds (85% frequency of seeing) were measured at 1-2 degrees eccentricity for one and two cone conditions following established procedures [20]. Experiments began by collecting a high SNR image from an average of 60-90 frames. Care was taken to select a region of the retina that would subsequently be used in appearance judgments. The experimenter then selected the center pixel of 8-12 contiguous cones from the reference image for testing. Thresholds were measured with an adaptive staircase procedure (QUEST) [35]. Each spot of light was monochromatic 543 nm, 0.35 arcmin (or 3×3 pixels) and was raster scanned against a low-photopic white background (40 cd/m^2^). In the case of paired stimulation, two spots of light, each 0.35×0.35 arcmin were delivered on each stimulus frame. Stimuli were presented over 500 ms (15 frames). The subject initiated each trial with a button press. An auditory beep indicated the start of the trial and then a stimulus was delivered to the center of either one or two of the selected cones. The subject reported whether she saw the flash with a single yes/no button press. No feedback was given.

Each session consisted of four interleaved staircases. Two staircases measured single cone thresholds and two measured paired stimulation thresholds. Each staircase terminated after 35 trials. Stimulus order was randomized. On each trial one cone or one pair from the pre-selected group was targeted. Therefore, these measurements reflected an average threshold over the 8-12 cones. Thresholds for specific cones or pairs could not be estimated from this data, since each location was only targeted on a handful of trials. This approach was an efficient way to approximate thresholds over a larger group of cones and allowed us to proceed more quickly to appearance measurements, which were our primary interest.

A fraction of the 8-12 cones selected at the start of the experimental session were separated by multiple cones. Variable distances between cones in the selected region was a potentially confounding factor. To minimize its effect, threshold measurements were only made between pairs of cones separated by no more than one cone or roughly two arcmin between the center of each cone. Cones at this eccentricity are about 1 arcmin in diameter. At the end of each session, threshold energy was estimated from each staircase using the QUEST mean procedure [36]. This generated four threshold estimates: two for single cones and two for pairs. We then averaged thresholds within each condition and compared the threshold energy between two- and one-cone conditions.

### Appearance judgments

Stimulus conditions in the appearance task were identical to the detection task. Flashes were 543 nm and 500 ms in duration and there was a low photopic white background. Each spot of light was 0.35 arcmin. Experimental sessions began by capturing a high SNR image of the subject’s cone mosaic. From that image, three contiguous cones were selected for study. By selecting contiguous cones, we assured that cones were never separated by more than one cone (a center-to-center distance of ~2 arcmin), which was the limit we set in the detection task. The subject initiated each trial with a button press, which was accompanied by an audible beep. On each trial, one or two of the selected cones were stimulated. The light energies used for one and two cone stimuli were set to each subjects’ previously determined detection thresholds. The recorded frequency of seeing in this task was 85.5%, as expected. Each cone and pair was tested 12 times for a total of 72 trials per session ([3 cones + 3 pairs] × 12 trials). Trials were randomly interleaved.

After each trial, subjects judged the hue and saturation with a scaling procedure [21, 37]. The subject indicated the percent of red, green, blue, yellow and white contained in each stimulus using five button presses such that each press represented 20% (5×20%=100%). This response scheme is called five category scaling. One subject, S20075, used an alternative response schemed, called 4+1 category scaling [37]. In this procedure, the subject first rated saturation on a seven point scale. Then, hue was rated with five button presses using only red, green, blue and white. It has been shown previously that these two procedures produce very similar results, but some subjects prefer the 4+1 category approach [37]. Both results were converted into a common metric space as described below.

### Color appearance analyses

The raw color appearance dataset contained a total of 4,968 trials completed by three subjects. Before analyzing the data, unusable trials were removed. The location of the stimulus on each frame was recorded in real-time with a digital cross written into the video frames. To identify unusable trials, a delivery error was computed as the standard deviation of the stimulus location over the 15 frames (500 ms). Trials with delivery error greater than 0.35 or less than 0.01 arcmin (values below 0.01 do not occur naturally) were considered unusable. In those trials, we could not be confident that the correct cone was targeted. After removing bad trials (3.6%), 4,788 trials remained for further analysis. The remaining trials had a mean delivery error of 0.19 arcmin (standard deviation = 0.036 arcmin), which was about 1/5 of the diameter of cones at the eccentricities tested. Trials that either targeted an S-cone or were not detected were also removed. The remaining dataset contained trials in which individual or pairs of L-and M-cones were stimulated (N=4,057). Finally, cones and pairs which had fewer than four good trials were not analyzed due to low statistical power. Most cones/pairs (71%) had at least 10 good trials.

Raw scaling data was transformed into a uniform appearance diagram [37]. For each trial, the number of red, green, blue, yellow and white button presses were converted to a percentage of the total button presses (five). A green-red dimension was computed as *gr* = (*green*% − *red*%)/100% and a yellow-blue dimension as *yb* = (*yellow*% − *blue*%)/100%. Saturation was computed from a sum of the absolute values of the green-red (gr) and yellow-blue (yb) dimensions (| *yb* | + | *gr* |). In 4+1 category scaling, each color category was scaled by the saturation judgment, which was normalized to range from 0-1. For example, consider a spot that was rated 60% red and 40% yellow at 40% saturation. Red and yellow, in this case, would be scaled down to 24% and 16%, respectively.

Analyses were carried out in the R programming language (https://www.r-project.org/).

### Data availability

The data and source code underlying the results presented in the study are available from GitHub at https://github.com/bps10/Schmidt-Boehm-Tuten-Roorda_2019.

## Results

### Detection thresholds sum linearly

The goal of these experiments was to determine how the visual system combines information across cones when making color judgments. To investigate this question, we probed cones individually or in pairs with an AOSLO microstimulator (543 nm; 500 ms; 0.35 arcmin). All tested cones were 1-2 degrees from the fovea. Before quantifying appearance, we first measured detection thresholds in the one and two cone conditions in order to control for differences in sensitivity. Threshold energy (threshold intensity multiplied by stimulus area) for achieving 85% frequency of seeing (FoS) was determined with an adaptive staircase procedure. Each subject completed at least four sessions of the detection task and each session contained two staircases per condition (one and two cone stimuli). At the end of the session, we averaged the threshold estimates for each condition and compared them. The values reported in Table 1 are the ratio of two:one cone threshold energies. This ratio equals one when the same energy (*i.e*. number of photons) was required to achieve threshold in both conditions. Values below one indicate less energy was necessary in the two cone case to achieve 85% FoS. The results from our three subjects were all close to one, which means, at threshold, each cone in a pair received approximately half the photons of the one cone case. Thus, the total energy was roughly equal across conditions and was consistent with linear summation. In subsequent experiments, individual and pairs of cones were stimulated at these threshold energies. Therefore, color judgments were made under conditions in which detection mechanisms were equally sensitive to all stimuli.

**Table 1.** Two:one cone threshold energy ratios

| subject | N | mean | StDv |
| --- | --- | --- | --- |
| S10001 | 4 | 0.98 | 0.09 |
| S20075 | 4 | 1.05 | 0.12 |
| S20076 | 11 | 0.92 | 0.1 |

N = the number of sessions completed. Each session contained two staircases for single cone stimulation and two for paired stimulation. After each session, the mean threshold energy in each condition was computed. The table reports the grand mean and standard deviation (StDv) of two:one cone ratios across all sessions.

### Variability in color perception

We next quantified color appearance of one and two cone spots presented at the measured threshold. Otherwise, stimuli were identical to those presented in the detection task. Three cones were selected for study in each session (Fig 1A). On each trial, either a single cone or a pair was targeted. After each flash, the subject judged the color of the spot using a hue and saturation scaling paradigm [21, 37]. Each cone and pair was tested twelve times. A total of 198 pairs were tested across three subjects. Hue and saturation scaling data were transformed into a color opponent representation. For each trial, the degree of perceived greenness versus redness and yellowness versus blueness was computed from percentage ratings as follows: *gr* = (*green*% − *red*%)/100% and *yb* = (*yellow*% − *blue*%)/100%. In this representation, saturation is expressed as the distance from the origin (in city block metric). A maximally saturated report falls along the outer diamond and a pure white response falls at the origin.

**Fig 1.**
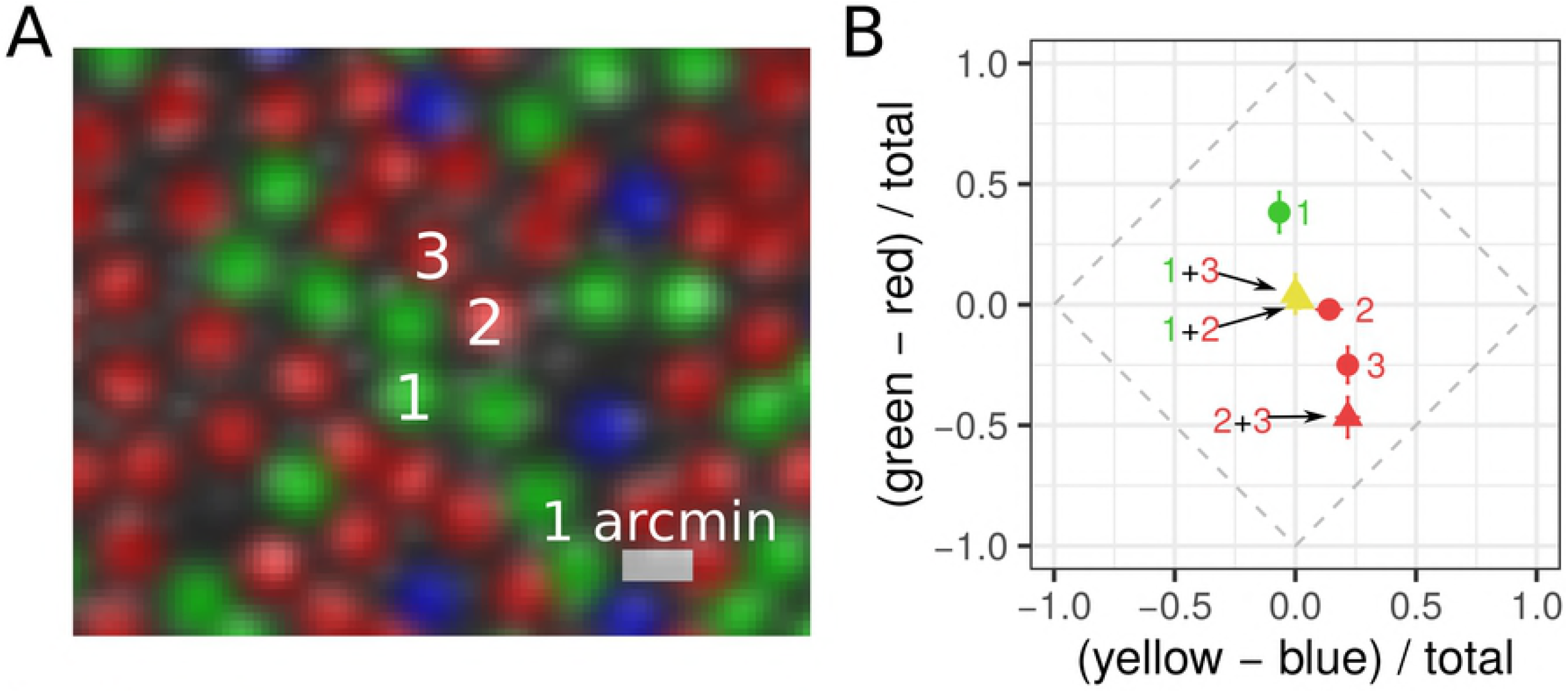
Measuring color appearance in one and two cone conditions. (A) Example AOSLO cone selection image (S20076; 1.5 degrees eccentricity). Groups of three cones were targeted during each experimental session (543 nm; 500 ms). Cones have been pseudo-colored to reflect their spectral type (red=L-cones, green=M-cones, blue=S-cones). The smaller, gray-scale blobs in between cones are rod photoreceptors. (B) Mean hue and saturation reports for one (circles) and two-cone (triangles) conditions. Numbers correspond to labels in (A). Results are plotted in a uniform appearance diagram (UAD), which represents bias towards the primary hues. An unbiased, or pure white, response falls at the origin. Green=M-cone(s), red=L-cone(s), yellow=L+M-cone pair. Error bars indicate ± SEM.

The results of one session are plotted in Fig 1B. In this example, Cone 1 was an M-cone and had a bias towards green (positive *gr* value). Cone 2 was an L-cone and elicited predominantly white reports. Cone 3, also an L-cone, was rated reddish-yellow (orange) with medium saturation (negative *gr* value, positive *yb* value). The percepts elicited when these cones were stimulated in tandem may provide insights into how the visual system combines color information across photoreceptors. In the example, when Cone 1 was targeted together with either Cone 2 or Cone 3, the average report had no clear color bias. In comparison, when Cone 2 and 3 were targeted they elicited a medium saturated orange report. Below, we analyze the results from all sessions and subjects.

We first grouped each trial based on which cone or pair was probed. The results are reported in Fig 2A and separated by subject. Each point in these plots represents the mean response measured from a single cone or pair. This plot illustrates the variability in responses across cones/pairs and between subjects. There are a few features to note. Firstly, there were individual differences in color responses: S20075 used blue more frequently than the two other subjects and S10001 did not report yellow on any trials. However, the general patterns are similar. Most of the variance was found along the green-red dimension and there were few points that fell in the blueish-red or greenish-yellow quadrants. Secondly, in two subjects with classified mosaics, we found L-cone targeted trials were red biased, while M-cones were green biased. These patterns were similar to previous reports from single-cone [21–23] and large-field studies [38]. Thirdly, within a single subject, there was considerable variability between cones and pairs with the same spectral sensitivity. Similar variability in single cone mediated percepts has been reported previously [21–24]. This is the first report of variability in percepts elicited from pairs of cones.

**Fig 2.**
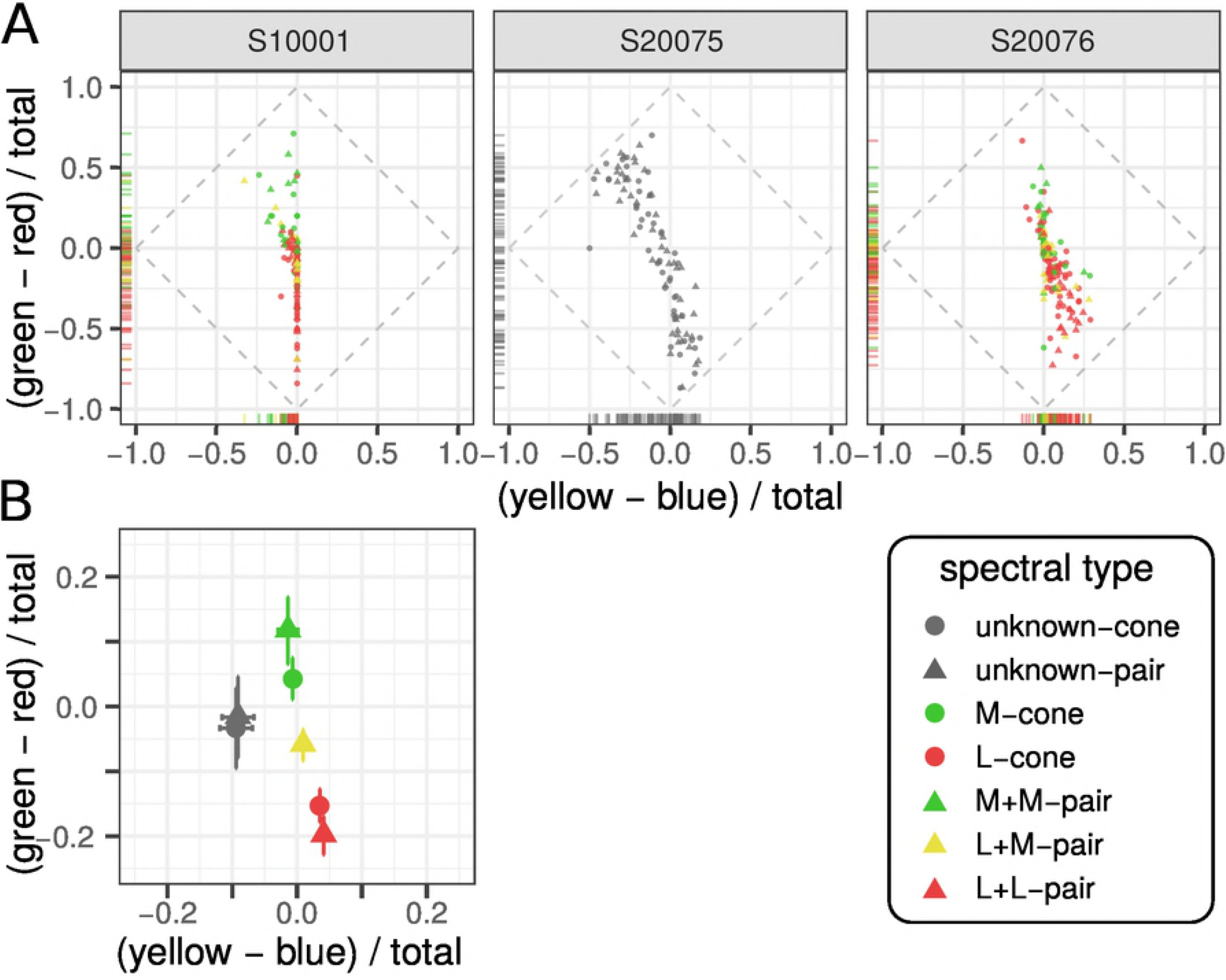
Number and type of cones probed influence color reports. (A) Average response from each cone and pair targeted in three subjects. Data was transformed into an opponent representation: yellow-blue and green-red. Marginal distributions are represented along each axis with rug plots. (B) The data from each subject was further grouped according to the cone type tested. The mean and standard error of each group are represented.

To better appreciate the influence of cone type and number of cones targeted on color reports, data was pooled across subjects and grouped according to the type of cone or pair probed. The mean and standard error for each group is shown in Fig 2B. When an individual or pair of M-cones was targeted the average *gr* response was greater than zero, indicating a bias towards green. In comparison, the average L-cone(s) elicited biases towards red and yellow. Together, these cone type specific differences in color reports were consistent with a predictive relationship between cone type and color report, as previously reported [21, 33]. Two cones with the same photopigment tended to elicit slightly more saturated reports than single cone trials. On the other hand, one L- and one M-cone targeted together tended to produce desaturated reports.

### Mosaic parameters do not predict percepts

Fig 2A illustrates that color reports varied even between cones with the same photopigment. Some L-cones, for instance, elicited saturated red percepts, while a majority produced white or desaturated red reports. We next asked whether this variability could be explained by features of the mosaic. Specifically, can we predict whether an L-cone will produce a saturated or a desaturated red based on the surrounding cone types? And in the case of paired stimulation, did the distance between the two cones influence color appearance? The existence of such relationships could implicate low-level neural mechanisms, such as chromatically-opponent ganglion cells, in this behavior.

The local neighborhood surrounding a cone is thought to be an important factor influencing color percepts associated with small spots [39]. To test this prediction, we found the number of L-cones in the immediate neighborhood of each cone/pair. In keeping with prior work [21–23, 40] the local neighborhood was defined as the six nearest cones. In the case of a pair, the immediate neighborhood for each cone was found separately and duplicates were removed. We did not find a significant correlation between the number of neighboring L-cones and the mean response in any dimension (*gr, yb* or saturation)(*p* > 0.05).

The distance separating two cones in a pair may also be an important factor influencing appearance. However, this measure was not correlated with hue or saturation reports (*p* > 0.05). Cone pairs were never separated by more than one cone, which may explain why we did not detect a relationship. Moreover, subjects verbally reported that the flashes always appeared as a single uniformly colored dot. In the future, systematically varying the distance between stimulated pairs will be an informative exercise. At a certain critical distance, the spots of light will be seen as two spatially distinct dots. It is less clear at what distance the spots will be perceived as two distinct colors.

### Paired simulation was predicted by an average of individual reports

While features of the mosaic and the physical stimulus did not predict color reports, we hypothesized that the responses recorded following individual cone trials would be predictive of paired stimulation. To assess this prediction, we matched the mean response from each cone pair with the the mean report from each cone tested individually. We then fit an averaging model to the data. Behavioral reports, *r*, from two-cone stimulation were predicted by an average of the individual responses: *r*_12_ = (*r*_1_ + *r*_2_)/2. Predictions were computed for *gr* and *yb* dimensions separately. Fig 3 shows the measured responses plotted against these predictions. An average of the single cone responses was a good fit to the data in both the *gr* (R^2^ = 0.73; *p* < 0.01) and *yb* (R^2^ = 0.75; *p* < 0.01) dimensions. The best fit lines had slopes close to unity, which further supported the conclusion that an average of individual responses was a good model.

**Fig 3.**
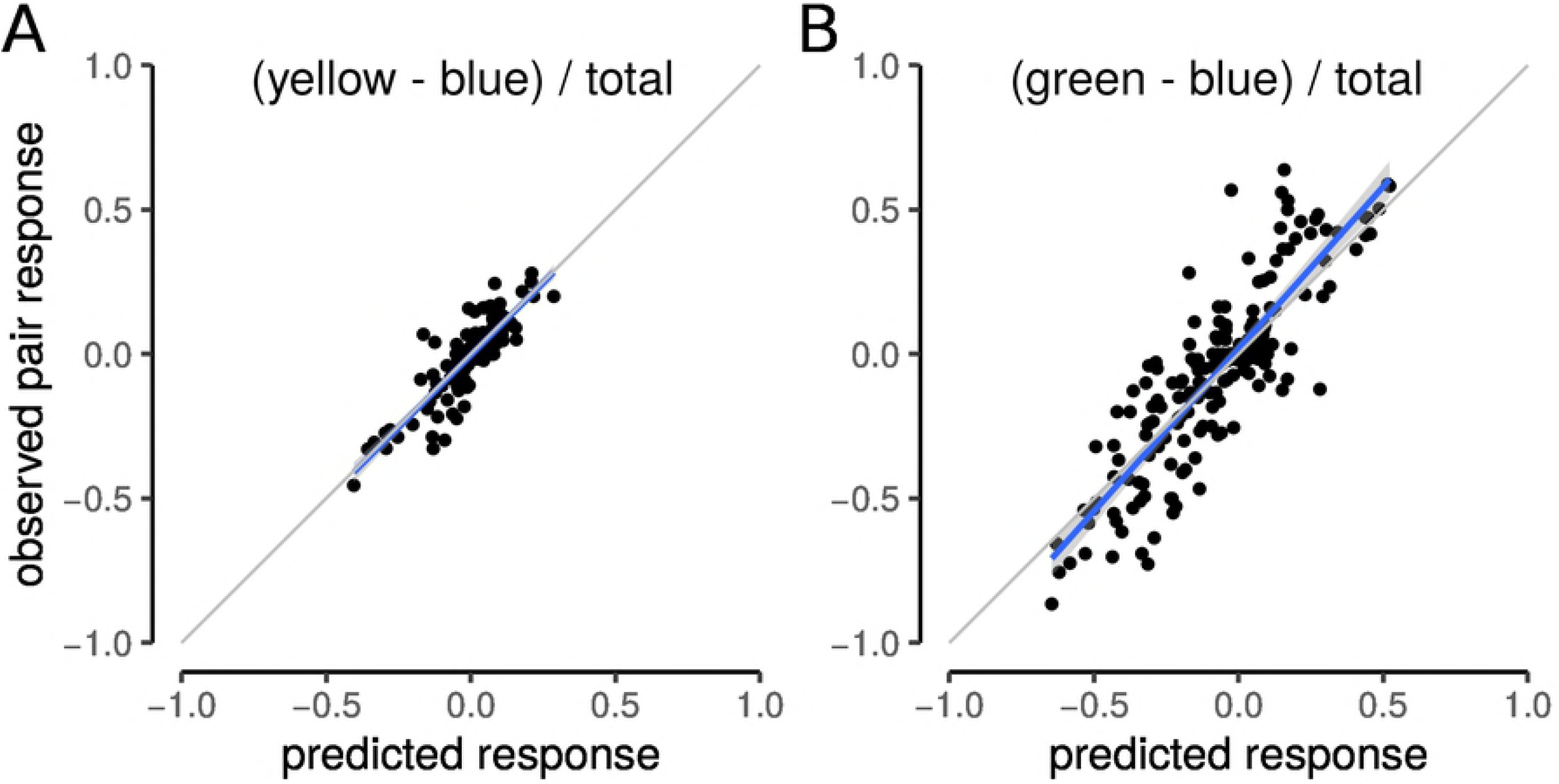
An average of individual responses predicts paired stimulation. The response measured for each pair was predicted from the average response of the two cones tested individually. Blue line represents the best fit line with 95% confidence intervals indicated by gray shading. (A) Blue-yellow (*by*) dimension. Best-fit line: *yb_observed_* = 0.004 + 1.002*yb_predicted_*. (B) Green-red (*gr*) dimension. Best-fit line: *gr_observed_* = 0.02 + 1. 12*gr_predicted_*. Gray line indicate unity slope.

### Super-saturation from cone-pairs with the same photopigment

While an averaging model captured a large fraction of the variance in two-cone color judgments, there were some pairs that deviated substantially from the best fit line. We wondered whether these deviations from an average might be predicted by the sub-class of the two cones. For instance, were L+M-pairs more likely to deviate from the model? To answer this question, we found the saturation for each pair and compared it to the saturation predicted by the average of the two cones probed alone (Fig 4A). A unity line represents the condition where the observed saturation judgment was predicted exactly by an average of individual responses. Notice that the L+L and M+M pairs tended to lie above the unity line, particularly at higher saturation values. In contrast, the L+M pairs often fell below the line. These observations indicate that cones of the spectral type produced slightly more saturated reports than predicted by the average of their individual responses.

**Fig 4.**
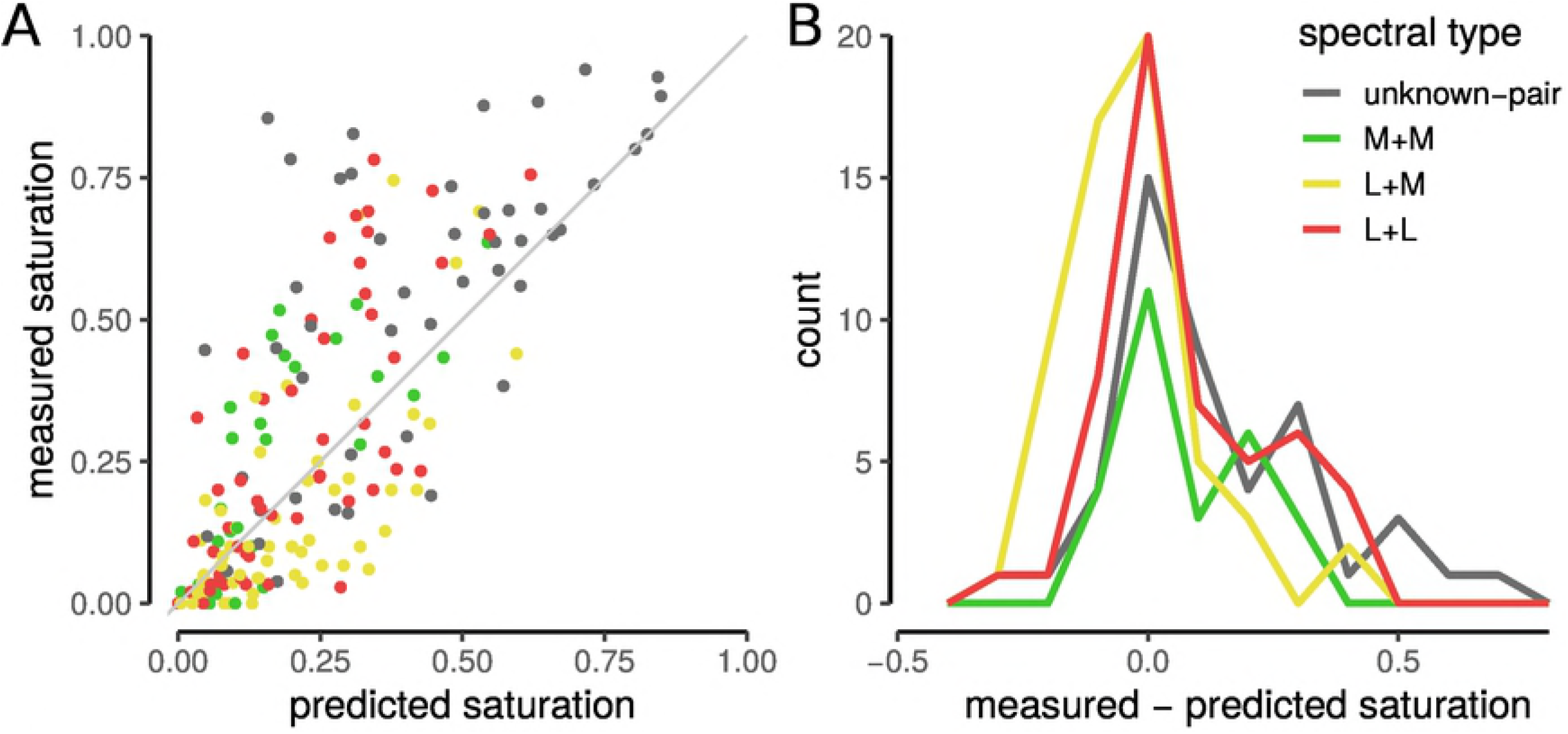
Cone pairs with the same spectral sensitivity produce higher saturation ratings than predicted. Saturation judgments were predicted for each measured cone pair with a linear average model. (A) Model predictions were plotted against the mean saturation ratings measured for each pair. Gray line indicates a prediction that matches the measured judgment exactly. (B) Distribution of measured responses minus predicted responses. Colors indicate cone type of pair: red=L+L, green=M+M, yellow=L+M, gray=unknown.

We quantified this trend directly by taking the difference between the observed and predicted saturation judgments. The results are illustrated in a histogram (Fig 4B). Student’s t-test’s confirm that the L+L and M+M pairs were significantly more saturated than an average of their individual responses (mean=0.072, *t*_78_=4.2, *p* < 0.01). In comparison, the mean difference for L+M pairs approached significance in the opposite direction (mean=-0.034, *t*_56_=-1.9, *p* = 0.06). These pairs were slightly more likely to be less saturated than predicted by an average of the individual responses. Across all of the unclassified cones tested in S20075, the average pair was more saturated than an averaging model predicted (mean=0.123, *t*_46_=4.0, *p* < 0.01). It is worth noting that this dataset contains all combinations of L and M cones. Had L+M pairs been removed from the S20075 dataset, the difference between observed and predicted saturation judgments may have been even more pronounced.

## Discussion

Small spots of light were targeted to individual or pairs of cones and color appearance was quantified. We report here that both the number and spectral type of targeted cones influenced color percepts (Fig 2). Pairs of cones elicited percepts that were predicted by an average of individual responses (Fig 3). When cones of different sensitivity (L+M) were targeted, their responses tended to cancel, or create a desaturated percept. When two cones from the same subclass were probed, their responses were generally consistent with their spectral sensitivity: on average, two L-cones appeared red and two M-cones appeared green. These findings were consistent with the involvement of a spectrally-opponent mechanism(s) [41–43]. The most surprising finding of the present work was that when two cone of the same type were probed, subjects reported seeing a hue that was more saturated than an average of the two probed alone (Fig 4). This was unexpected because the stimuli were adjusted to be equally detectable (Table 1). Thus, while a detection mechanism was equated across conditions, a second, color sensitive mechanism, was influenced by activity in a second cone and saturation was systematically elevated.

Why would two cones produce a more saturated percept when targeted together? The visual system generates color by comparing relative activity in the three cone classes. For example, a green surface excites M-cones more than L- or S-cones. Activation of a single M-cone also elevates the activity in M-cones relative to L+S and thus may appear green. However, activity in a single M-cone could equally well be caused by a broadband (white) spot of light, because individual cones are sensitive to changes in wavelength and intensity [44]. Thus, small spots, like the ones presented here, force the visual system to select, from multiple possible interpretations, the physical stimulus that most likely generated the incoming pattern of activity. This uncertainty was likely one reason why single M-cone trials sometimes looked white and sometimes green (Fig 2). The finding that even cones with the same photopigment elicited different percepts suggests that the visual system uses prior experience when making this guess [21, 25]. Stimuli not confined to one or two cones, for example larger stimuli or small spots which are allowed to move across the retina in a manner consistent with fixational eye motion, should provide the visual system with more information about the stimulus. Thus, the number of possible interpretations by the higher level visual system decreases. We hypothesize that when two M-cones were activated with an equally intense light the balance was tipped towards the interpretation that the physical stimulus was a uniform, middle-wavelength surface. In other words, activity in a second M-cone slightly decreased the likelihood that the physical stimulus was a broadband (achromatic) spot. Together, these observations suggests that the visual system uses a combination of spectral opponency and prior experience to assign color to small spots.

One possible mechanistic explanation for the increased saturation we observed in two cone stimulation (Fig 4) is the presence of a saturating non-linearity before cone signals are summed. Horwitz and Hass [45] described color cells in primary visual cortex that compressed cone inputs before summation in a manner consistent with our observations. In comparison, our threshold measurements followed a linear summation model, which was consistent with the area of complete summation (Ricco’s area) at this eccentricity [5]. Together, our observations support the idea that separate neural mechanisms mediated these two tasks [46], potentially implicating the parvo- and magnocellular pathways, which are thought to provide the basis for detection of chromatic and luminance differences, respectively [47].

There were a few factors potentially confounding the present work. Firstly, different stimulus intensities were used in one- and two-cone conditions. In order to equalize detection, each cone in a pair was stimulated with about half the intensity of single cone trials. An implicit assumption in the averaging model (Fig 3) was that hue and saturation judgments were not influenced by intensity. Our findings would require a more complicated model if this assumption was invalid. We have previously found that color judgments in single cone conditions are approximately constant over the range of intensities used in this study [21]. Therefore, we do not believe that stimulus intensity influenced our results.

Secondly, we cannot rule out the possibility that two-cone appearance judgments were influenced by a weighting mechanism at the site of spatial pooling [48]. However, in our study, two cone appearance judgments were well predicted by a simple average of their individual activations (Fig 3). Consequently, if the chromatic mechanism weighted cones prior to summation, those weights were small and had only a minimal effect on the measured responses. A final possible complication was that detection judgments were not made in a cone or pair-specific manner. We measured average cone/pair thresholds drawn from groups of 8-12 cones. Bruce [6] measured detection summation between cone pairs at a similar eccentricity and found that a subset of pairs followed a sub-linear summation rule. We cannot rule out the possibility that some of the pairs in our dataset also followed this strategy. However, Bruce’s [6] findings went in the opposite direction of our appearance results. A fraction of their cone pairs were less sensitive than either one tested alone. Thus, these small deviations cannot explain the increased saturation we found during paired stimulation trials.

The approach used here of targeting small groups of cones provides a means of testing more sophisticated hypotheses about early stage neural mechanisms and their role in shaping visual experience. Our evidence supports the idea that the appearance of small spots is dependent upon both the number and type of cones targeted. These observations are consistent with different strategies for combining information within versus across neuronal sub-classes. In the future, scaling these experiments to larger groups of cones will provide important clues about how the visual system extracts color and spatial signals in more naturalistic settings.

## Acknowledgments

We are grateful for technical assistance from Pavan Tiruveedhula. This work was supported by grants from National Eye Institute National Institute of Health awarded to A.R. (R01EY023591), B.P.S. (F32EY027637) and A.E.B. (T32EY7043-38). A.E.B. was also supported by the Minnie Flaura Turner Memorial Fund for Impaired Vision Research and the Michael G. Harris Ezell Fellowship.

## Competing interests

A.R. has a patent (USPTO#7118216) assigned to the University of Houston and the University of Rochester which is currently licensed to Boston Micromachines Corp (Watertown, MA, USA). Both he and the company stand to gain financially from the publication of these results.

